# Multiple pathways of LRRK2-G2019S / Rab10 interaction in dopaminergic neurons

**DOI:** 10.1101/2020.09.28.316992

**Authors:** Alison Fellgett, C. Adam Middleton, Jack Munns, Chris Ugbode, David Jaciuch, Laurence Wilson, Sangeeta Chawla, Christopher J. H. Elliott

**Affiliations:** Department of Biology, University of York, York, YO1 5DD, UK; Department of Physics, University of York, York, YO1 5DD, UK; York Biomedical Research Institute, Department of Biology, University of York, YO1 5DD, UK

**Keywords:** *Drosophila*, dopamine, Leucine-rich-repeat-kinase2, bradykinesia, vision, circadian rhythms, sleep, courtship memory

## Abstract

**Background:** Inherited mutations in the LRRK2 protein are the most common known cause of Parkinson’s, but the mechanisms by which increased kinase activity of mutant LRRK2 leads to pathological events remain to be determined. *In vitro* assays (heterologous cell culture, phospho-protein mass spectrometry) suggest that several Rab proteins might be directly phosphorylated by *LRRK2-G2019S*. Which Rabs interact with LRRK2 in dopaminergic neurons to facilitate normal and pathological physiological responses remains to be determined. An *in vivo* screen of Rab expression in dopaminergic neurons in young adult Drosophila demonstrated a strong genetic interaction between LRRK2-*G2019S* and Rab10. We now ask if Rab10 is required for LRRK2-induced physiological responses in DA neurons.

**Methods:** *LRRK2*-*G2019S* was expressed in Drosophila dopaminergic neurons and the effects of Rab10 depletion on Proboscis Extension, vision, circadian activity pattern and courtship memory determined in aged flies.

**Results:** Rab10 loss-of-function rescued bradykinesia of the Proboscis Extension Response (PER) and visual defects of aged flies expressing LRRK2-G2019S in DA neurons. Rab10 knock-down however, did not rescue the marked sleep phenotype which results from dopaminergic expression of *LRRK2*-*G2019S*. Courtship memory is not affected by LRRK2 expression, but is markedly improved by Rab10 depletion. Anatomically, both LRRK2-G2019S and Rab10 are seen in the cytoplasm and at the synaptic endings of dopaminergic neurons.

**Conclusions:** We conclude that, in Drosophila dopaminergic neurons, Rab10 is involved differentially in LRRK2-induced behavioral deficits. Therefore, variations in Rab expression may contribute to susceptibility of different dopaminergic nuclei to neurodegeneration seen in people with Parkinson’s.

**Graphical Abstract:** Rab10 depletion ameliorates the proboscis extension bradykinesia and loss of synaptic signalling in the retina induced by *LRRK2*-*G2019S* expression (magenta arrows / orange crosses). Rab10 manipulation does not affect the ‘sleep’ phenotype from *LRRK2*-*G2019S* (magenta arrow). Reduction of Rab10 facilitates conditioned courtship memory, but LRRK2 has no effect (yellow arrow). All manipulations of Rab10 and *G2019S* in dopaminergic neurons, shown in the outline of the brain (filled cells have high levels of Rab10). We conclude that Rab10 and LRRK2 interact in some, but not all dopaminergic neurons. This may underlie differences in the susceptibility of different human striatal dopaminergic cells to Parkinson’s and explain why different symptoms initiate particular ages.

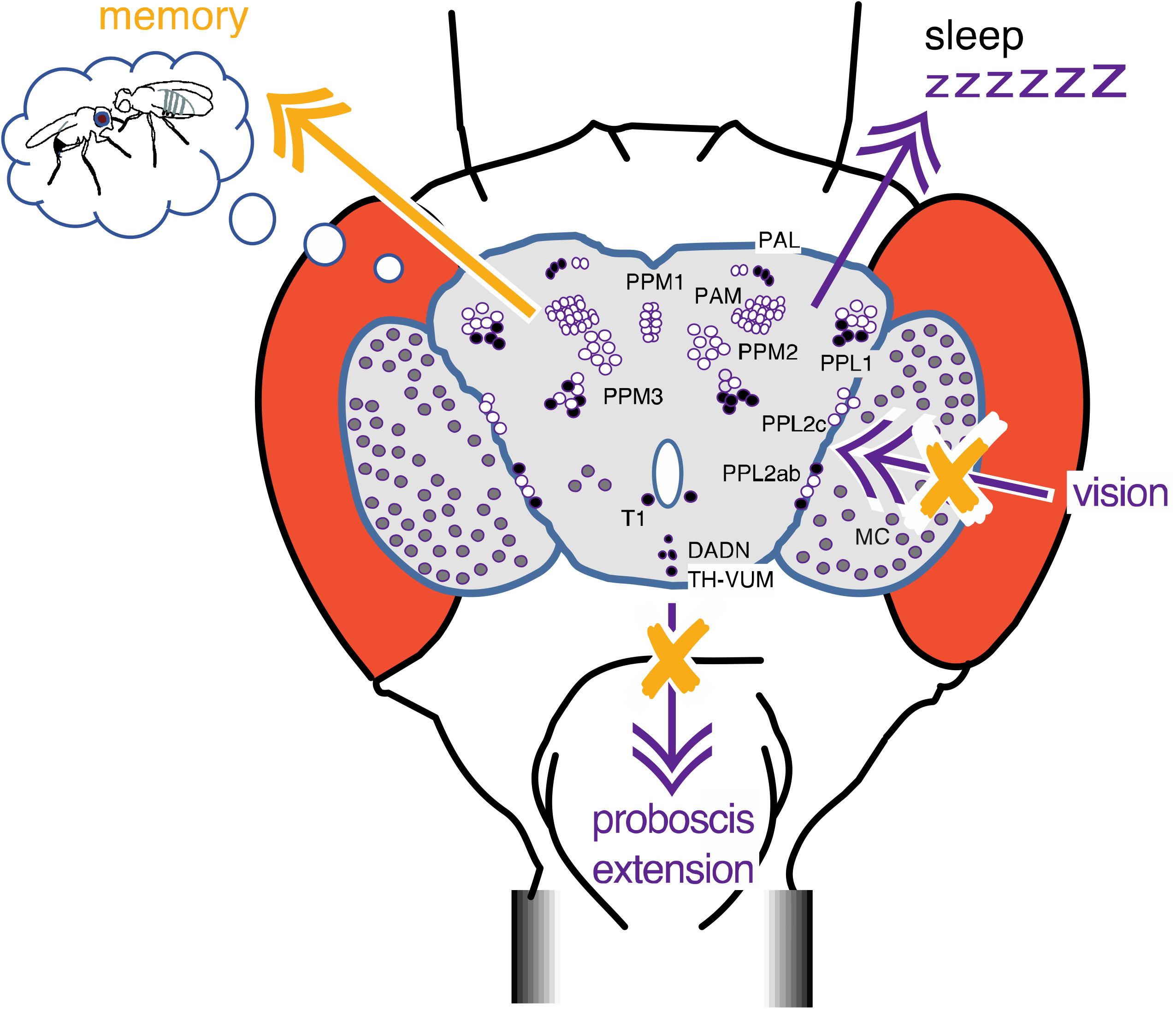

## Background

Mutations in the *LRRK2* gene are the most frequent genetic cause of late onset Parkinson’s. The *G2019S* mutation increases LRRK2 kinase activity [1], leading to a toxic cascade that kills dopaminergic neurons in the *substantia nigra*. This results in bradykinesia and sleep disturbances, while loss of dopaminergic amacrine cells in the retina contributes to visual deficits. However, the main steps between LRRK2 and these pathological outcomes remain to be determined. One of the first steps that has been suggested is that LRRK2-G2019S may phosphorylate a number of small GTPases [3,5,8,10,29,35 & 43] [2]. Cell culture experiments have shown a particular effect on Rab10 [2–7], (see [8] for review), while a visual screen of Drosophila synaptic processing indicated a particular synergy between Rab10 and G2019S [9]. As a molecular switch, phosphorylation of a Rab protein could result in changed effector binding with a series of downstream consequences. However, the level of Rab10 is quite disparate, even among different types of dopaminergic neurons [9,10] so that LRRK2 might not work in the same way in all dopaminergic neurons.

Since most genetic mouse models of Parkinson’s have very weak phenotypes, with lack of neuropathology [11], we turned to the fly where manipulation of Parkinson’s disease related genes leads to marked phenotypes: bradykinesia [12], loss of dopaminergic neurons [13], tremor [14], retinal degeneration [15] and sleep disorder [16] (see, for review [17]). As in mammals, fly dopaminergic neurons regulate movement, vision, sleep and memory.

To determine how important Rab10 is in the pathological cascade initiated by LRRK2 activity, we now test the knock-down / knock-out of Rab10 in dopaminergic neurons in the intact organism. We find the movement and visual deficits seen in flies expressing *LRRK2*-*G2019S* in dopaminergic neurons are rescued by the depletion of Rab10. However, the marked sleep phenotype caused by dopaminergic expression of LRRK2-G2019S is not affected by manipulation of Rab10. Conditioned courtship memory is not affected by LRRK2, though dopaminergic Rab10 reduction markedly improves this. We therefore find a role for Rab10 downstream of activated LRRK2 in some, but not all, dopaminergic neurons in the activation of LRRK2-associated physiological deficits in Drosophila.

## Materials and Methods

### Flies

All flies tested were male *Drosophila melanogaster*, using *TH*-GAL4 (kind gift of Serge Birman) to manipulate dopaminergic neurons or *nSyb*-Gal4 (from Julie Simpson (corresponding Bloomington stock 51635) for pan-neuronal expression. *Gr5a*-LexA was used to express the LexOp-*ReachR* channelrhodopsin in the sugar sensitive neurons independently from the GAL4-UAS manipulations. Flies were raised and crossed using standard Drosophila protocols [9]. The full list of fly lines is in Table 1.

### Validation of Rab10 knock-down and phosphorylation by *G2019S*

We used three strategies for Rab10 reduction. (i) We used a CRISPR /Cas9-generated *Rab10* null [18], in which Rab10 was severely reduced (both pan-Rab10 and phospho-Rab10 to less than 5% of wild-type, Fig. 1). This line was not used in the visual assay because the retinal RFP marker causes the eyes to fluoresce and this would distort the disco-chamber data. (ii) We used *Rab10*^*RNAi*^. This depleted phosphoRab10 (Fig. 1), but a small amount (~15%) of Rab10 was still present (Fig. 1). (iii) We used the deGradFP technique to knockout *YFP-Rab10* protein with the *vhhGFP* nanobody, to target *YFP-Rab10* to the proteasome. This was used in flies in which the wild-type Rab10 had been replaced by homologous recombination so that the *YFP-Rab10* was expressed at wild-type levels. This reduced the amount of phospho-Rab10 to ~50% of wild-type (Fig. 1). In the *GFP-Rab10* background, expression of a RNAi against GFP depleted some phospho-Rab10, but not as much as the nanobody, so we did not use this further.

**Fig. 1.**
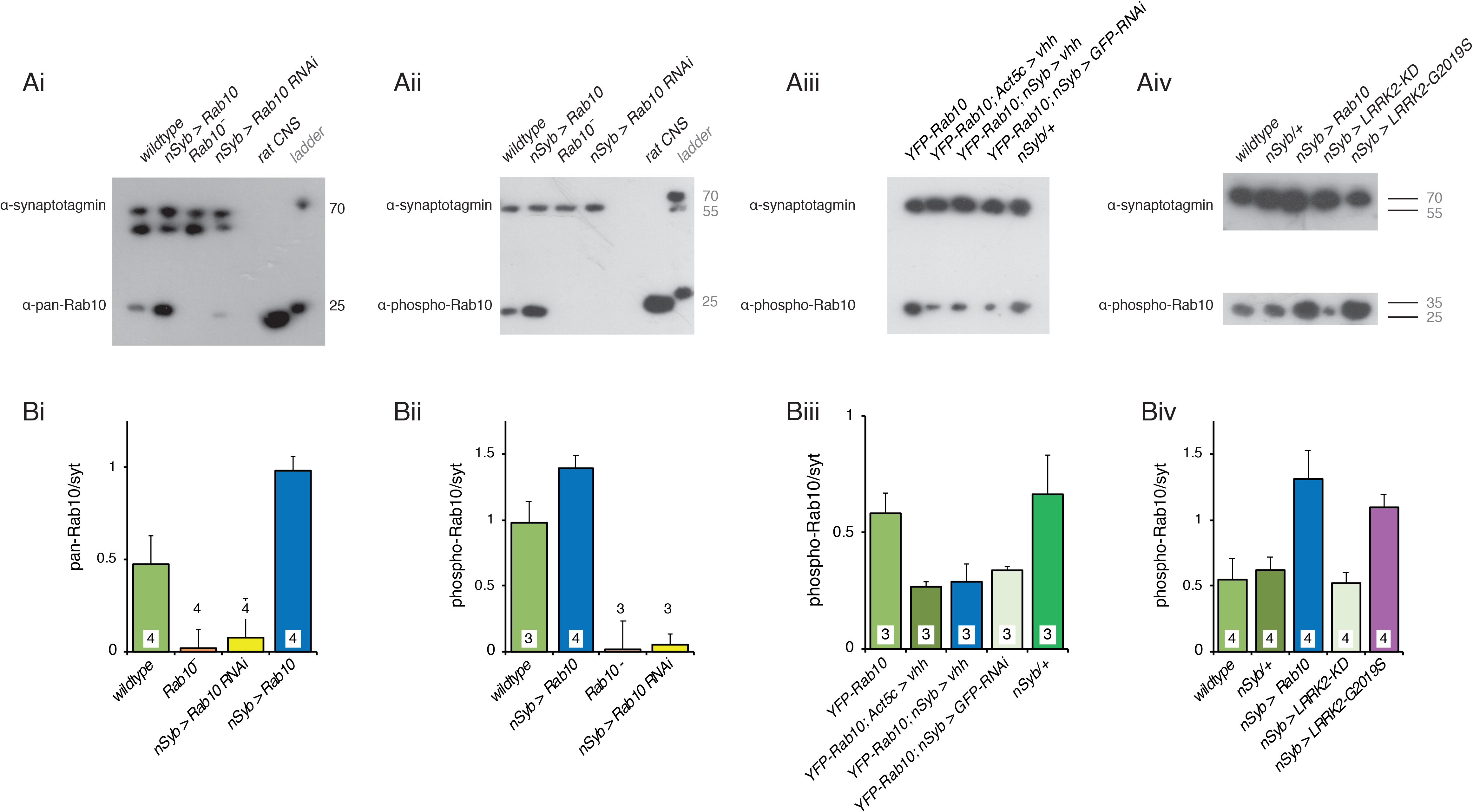
Validation of Rab10 depletion and phosphorylation *in vivo*. A. A pan-hRab10 antibody detects dRab10 and shows that Rab10 is increased by neuronal (nSyb-GAL4) expression of UAS-*Rab10*, abolished in the Rab10^−^ null, and substantially reduced by neuronal expression of *Rab10*^*RNAi*^. Antibody-specificity was confirmed by rat CNS binding. B. A phosopho-hRab10 antibody detects phospho-dRab10, which is increased by neuronal expression of UAS-*Rab10*, but undetectable with the Rab10^−^ null or with neuronal expression of *Rab10*^*RNAi*^. C. Validation of deGradFP technique. In flies where endogenous *Rab10* has been replaced by *YFP-Rab10*, global (Act5c-GAL4) or neuronal expression of the vhhGFP nanobody reduces the level of YFP-Rab10 by ~50%. A slightly smaller reduction is achieved by neuronal expression of *GFP*^*RNAi*^. D. Neuronal expression of *Rab10* increases pRab10, as does neuronal expression of *LRRK2*-*G2019S*. Neuronal expression of a kinase-dead *LRRK2* (*KD*, *LRRK2*-*G2019S-K1906M*) has no effect on the phosphorylation of Rab10. B. Quantification of the Western blots in the corresponding panels of A. Loading control: α-drosophila-synaptotagmin (α-syt). Exact genotypes and in Table S2.

The ability of LRRK2-G2019S to phosphorylate Rab10 *in vivo* was confirmed (Fig. 1Aiv; Biv), by comparing *nSyb > LRRK2*-*G2019S* with the kinase-dead (KD) form (*LRRK2-G2019S-K1906M*). In the *nSyb > LRRK2*-*G2019S* flies, the level of pRab10 was twice that of the controls, similar to the effect of pan-neuronal expression of *Rab10*.

### Western Blots

The levels of Rab10 and LRRK2-G2019S were assayed by Western Blot using standard protocols [9,14]. For Rab10 the antibodies were: α-pan-Rab10 (Nanotools, clone 605B11), α-phospho-Rab10 (Abcam, ab230261, 1:1000), and for LRRK2 (Neuromab, clone N241A /34). We used α-synaptotagmin 91 as a loading control [19]. Selectivity of Rab10 antibodies was confirmed by using a rat CNS extract [9]. For assay of *GFP-Rab10*, we used Guinea pig anti-GFP (Synaptic Systems, 1:1000) as described recently [9]. Quantification of the blots was carried out in Fiji.

**Bradykinesia** was assessed using an optogenetic stimulus. Flies were fed retinal (1 mM) pipetted onto the surface of their food for 1 week in the dark at 29 °C. They were restrained at 25 °C for 3 hours before the proboscis extension responses were observed with a Grasshopper 3 (Point Grey) camera mounted on a Zeiss Stemi microscope at 200 frames / second. A single flash was delivered from a ThorM470L3 LED, driven at 8 V for 7 ms. The stimulus was transcoded by LexOp-*ReachR* expressed in the Gr5a neurons. The area of the video occupied by the proboscis was automatically analysed by python code https://github.com/biol75/PER, and the Tukey - post-hoc test applied in R.

**Akinesia** was recorded from 1 week old flies, kept in the dark at 29 °C. Flies were restrained as described [14] and starved at 29 °C for 3 hours before being offered a droplet of 100 mM sucrose three times. Each response was scored Yes /No and the median response for each fly used. The χ^2^-post-hoc test (Fifer) was done in R (3.3.3).

### Visual assays

On the day of emergence, flies were placed in the dark or in disco-chambers at 29 °C. 1 week old flies were prepared for SSVEP (Steady State Visual Evoked Potential) measurements as recently described [9]. Stimuli were generated and responses recorded by an Arduino Due system with FFTs and contrast sensitivity computed in Matlab. Dunnett’s post-hoc test was applied in R. Full code at https://github.com/wadelab/flyCode. DPP (Deep-Pseudo-Pupil) images were captured with a Dino-Lite Camera and software, and images cropped, quantified and converted to grey scale in Fiji. Both images in Fig. 3Aii were combined in Fiji before the colour was removed and the contrast increased.

**Circadian rhythms and sleep** patterns were recorded as described recently [20] with a TriKinetics DAM system. Flies were placed in the monitor at 29 °C on the day of eclosion, and locomotor activity was recorded in 1 minute bins for 3 days under 12:12 h light /dark cycles followed by 7 to 10 days in constant darkness. Activity data was analysed using the ActogramJ plugin for ImageJ and sleep data using the ShinyR-DAM code [21] using the branch at https://karolcichewicz.shinyapps.io/ShinyR-DAM_3_1_Beta/.

**Conditioned courtship memory** was recorded using the well-developed conditioned courtship memory protocol for Drosophila [22,23]. One-week old males, (kept at 29 °C), were provided with a virgin female and the time spent in courting behaviours measured. Half of the males had recently (~ 30 minutes before) spent an hour with a mated female; half were naïve. Males who fail to ‘remember’ their rejection will spend more time courting the virgin female than controls. The Courtship Index (CI) was calculated as the percentage of ten minutes that a male spent courting; and the memory index (MI) by dividing the CI of each test fly by the mean CI of the naïve /sham flie s of the same genotype. A score of 0 indicates highest memory performance possible and a score of ≥1 indicates no memory. The distribution of MI was compared between genotypes using the Kolmogorov-Smirnov test. All naïve and trained groups contained 16-23 flies.

**Immunocytochemistry** was performed on 3-5 day old flies expressing *G2019S* as described recently [14], using the LRRK2 mouse antibody Neuromab (clone N241A /34). In some flies, mCD8-GFP was also expressed in the dopaminergic neurons. Each figure is representative of three preparations from at least two crosses.

## Results

To investigate a role for Rab10 in dopaminergic neurons, downstream of activated LRRK2, we recombined the *Tyrosine-Hydroxylase* GAL4 with UAS-*G2019S* and designated these ‘PD-mimic’ flies as *THG2*. We compared these flies with controls in four types of behavioral /physiological phenotypes: bradykinesia in the proboscis extension response, neural signalling in the retina, sleep patterns and conditioned courtship memory. These four systems are controlled by different clusters of dopaminergic neurons.

### Rab10 reduction rescues LRRK2-G2019S bradykinesia

The major features of Parkinson’s are movement deficits (bradykinesia), slowing or loss of movement and tremor. We therefore begin our analysis with the movement of PD-mimic *THG2* flies. The Proboscis Extension Response (PER), a reaching movement used by Dipteran flies to obtain food, is particularly amenable to analysis [24] (Fig. 2Ai). When a fly walks across a surface and encounters a droplet of sugary solution, the sucrose-sensitive neurons on the legs are stimulated and this elicits a rapid extension of the proboscis. The PER is modulated by a single dopaminergic neuron, the Tyrosine Hydoxylase Ventral Unpaired Median (TH-VUM) neuron [25], which has a high level of Rab10 expression [9]. Expression of *G2019S* in this neuron results in movement deficits – akinesia (loss of the PER), slower response (bradykinesia) and tremor [14]. Further, the PER can be elicited using an optogenetic stimulus. In this paradigm, the sucrose-sensitive sensory neurons in the leg express a channelrhodopsin which is stimulated by a flash of light. Expression of a channelrhodopsin by the LexA /LexOp system in the sugar-sensitive ‘Gr5a’ neurons on the leg is independent of the expression of *LRRK2*-*G2019S* in the TH-VUM and other dopaminergic neurons by the GAL4-UAS system [26]. This precise control of the stimulus makes it possible to measure the time course of the reaching movement, and separates the activation of the stimulus from manipulations using the GAL4 system in the dopaminergic neurons.

**Fig. 2.**
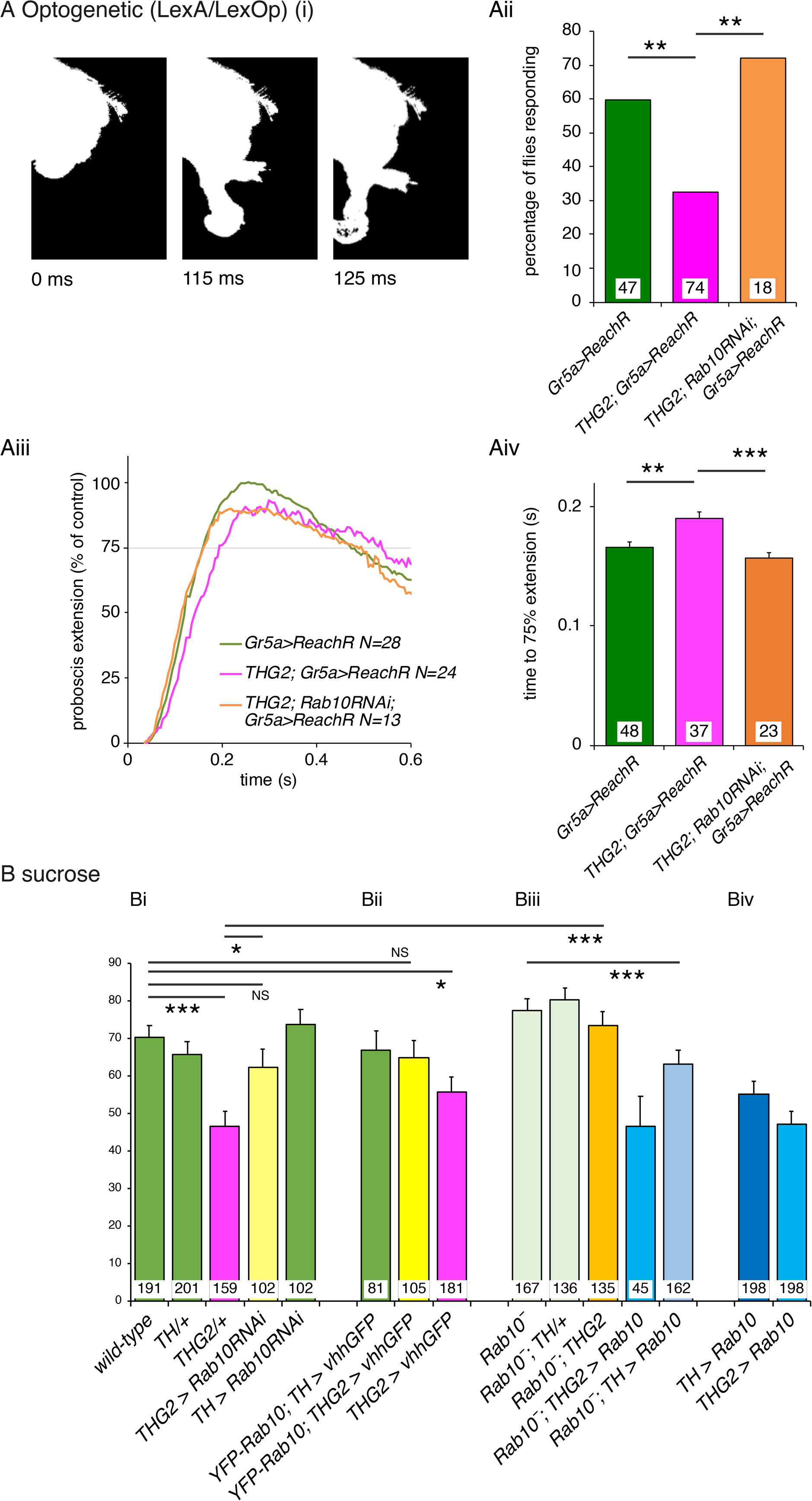
*Rab10* knock-down rescues *LRRK2*-*G2019S*-induced bradykinesia. **A. Optogenetic stimulation of the Proboscis Extension Response (PER).** Ai. To reach for food or liquid, the fly extends its proboscis in response to an optogenetic stimulus to sensory neurons on the legs. The full extension response is shown in Movie M1. Aii Expression of *LRRK2*-*G2019S* in dopaminergic neurons (*THG2*) reduces the proportion of flies that respond to a single flash of light, and this is rescued by Rab10 reduction with *Rab10RNAi*. Aiii. Dopaminergic reduction in Rab10 rescues the bradykinesia (slower response) of flies expressing *G2019S* in their dopaminergic neurons. Aiii. Raw traces; Aiv. mean data. To respond to the optogenetic stimuli all flies carry *LexA/Op Gr5a>ReachR*. **B. Sucrose stimulation of the PER**. Bi. Flies expressing *G2019S* in their dopaminergic neurons (*THG2*, magenta bars) respond less frequently to sucrose than wild-type flies (green) i.e. they show akinesia. This is rescued in *THG2* flies with dopaminergic reduction in Rab10 using *Rab10*^*RNAi*^ (*THG2* > *Rab10*^*RNAi*^). Bii Reduction of dopaminergic Rab10 with the deGradFP technique (*Rab10 GFP; THG2>vhhGFP,* yellow bars) also rescues the *G2019S*-induced akinesia. Biii. The Rab10 null (*Rab10^−^*, orange bar) also reverts the *G2019S* deficit, while expression of *Rab10* in the null background again induces akinesia (*Rab10^−^; TH > Rab10*, light blue bar). Biv. By itself, *Rab10* expression phenocopies *THG2*. Exact genotypes and full statistical data in Table S3.

In our optogenetic setup, 60% of control flies respond to a single flash of light by smoothly extending their proboscis, whereas flies in which *Tyrosine-Hydroxylase* GAL4 was used to express *LRRK2*-*G2019S* (*THG2*) only respond 30% of the time, i.e. *G2019S* induces akinesia (Fig. 2Aii). Those *THG2* flies that do respond, take ~100 ms longer to achieve maximum extension, showing bradykinesia (Fig. 2Aiii, iv). Knock-down of Rab10 using *Rab10*^*RNAi*^ co-expressed with *LRRK2*-*G2019S* in the dopaminergic neurons, fully reverts both akinesia and the bradykinesia (Fig. 2Aii-iv).

We next tested if Rab10 knock-down would also rescue the deficits induced by *G2019S* in response to the natural stimulus, a sugar solution. Just over two-thirds of control flies respond to sucrose (70%, Fig. 2Bi, green bars), whereas less than half of *THG2* flies respond to sucrose (46 %, Fig. 2Bi, magenta). This deficit is fully rescued when *Rab10*^*RNAi*^ is co-expressed with *THG2* (*THG2* > *Rab10*^*RNAi*^ Fig. 2Bi, yellow). Dopaminergic expression of *Rab10*^*RNAi*^ alone has no effect on the proportion of flies that respond to the sucrose solution (*TH > Rab10*^*RNAi*^, Fig. 2Bi, green).

As RNAi may have off-target effects, we supplemented this with a nanobody-mediated-protein knockdown technique, deGradFP [27,28]. In this, we deployed flies in which the wild-type *Rab10* gene had been replaced by *YFP-Rab10* by homologous recombination. Then we expressed an anti-GFP nanobody (vhhGFP) to deplete YFP-Rab10 protein in just the dopaminergic neurons using *TH*-GAL4. In this experiment, *YFP-Rab10* flies with both *G2019S* and *vhhGFP* expression (Fig. 2Bii, yellow bar) responded identically to wild-types, or *YFP-Rab10* controls in which dopaminergic neurons expressed just *vhhGFP* (Fig. 2Bii, *YFP-Rab10; TH > vhhGFP* green bar). This indicates a full rescue of the *THG2* induced akinesia. We confirmed that expressing *G2019S* and *vhhGFP* in flies with wild-type *Rab10* still resulted in akinesia (Fig. 2Bii, magenta bar).

While the *Rab10* knockout is lethal in mammals [29], the fly *Rab10* null (*Rab10*^−^) is viable [18]. We therefore tested if a global removal of Rab10 would also ameliorate the akinesia deficit in the *THG2* flies. In the *Rab10*^−^ background, the *THG2* Proboscis Extension Response is fully rescued (Fig. 2Biii, orange), to the same level as wild-type controls (Fig. 2Bi, green). In control experiments, the proportion of *Rab10*^−^ flies that responded to sucrose was identical to wild-types; this was true for the homozygote stock and for an outcross to *TH* in males (*Rab10* is on the X-chromosome)(Fig. 2Biii, pale green).

Rab10 is known to be phosphorylated by LRRK2 *in vitro* [2] and in the fly, *in vivo* (Fig. 1). Phosphorylation is likely to be part of the activation mechanisms of Rab10. If increasing phosphorylation of Rab10 is a key event in *LRRK2-G2019S* driven dysfunction in dopaminergic neurons, we would expect dopaminergic expression of *Rab10* to phenocopy that of *G2019S*. We found that 55% of such *TH > Rab10* flies responded to sucrose, in an almost identical manner to the *THG2* flies (Fig. 2Biv) with pronounced akinesia. No further increase in akinesia is seen when both *G2019S* and *Rab10* are expressed at the same time.

To confirm that akinesia was solely dependent on dopaminergic Rab10, we took the *Rab10* null, and crossed it with flies expressing Rab10 in just the dopaminergic neurons. Such *Rab10^−^; TH > Rab10* flies have akinesia when compared with the *Rab10^−^* flies (Fig. 2Biii). Similarly, the *Rab10^−^; THG2 > Rab10* show less response than *THG2* > *Rab10* flies.

Thus, the *G2019S*-induced movement deficits in the PER are rescued by depleting Rab10 in the dopamine neurons either at the mRNA level using RNAi, or at the protein level using a nanobody, or with the global null. Increasing Rab10 in the dopaminergic neurons induces akinesia similar to the expression of *LRRK2*-*G2019S*.

### Dopaminergic reduction in Rab10 rescues G2019S-induced visual deficits

People with Parkinson’s also show visual deficits including loss of dopaminergic neurons from the retina [30,31]. Aged *THG2* flies also show strong retinal degeneration with vacuoles throughout the optic lobe [15]. To demonstrate retinal degeneration, we deployed the ‘Deep-Pseudo-Pupil’ (DPP) assay [32]. When wild-type flies are illuminated from below, they show a DPP, with about 6-8 glowing ommatidia. In this, the normal retinal structure focuses light from directly below towards the observer through the center of the ommatidium, whereas light which is off-center is blocked by the pigment granules at the edge of the ommatidium (Fig. 3Ai). We tested several types of genetic background for the *THG2* ‘PD—mimic’ flies used in movement assays and found that the contrast between DPP and the red eye pigment was best when the *THG2* was in the deGradFP background. Although the DPP was clear in these young flies, in flies aged for 14 days, the DPP is distorted in these *THG2;vhhGFP* flies (Fig. 3A ii-iv). The outline of the DPP is irregular, rather than a neat ellipse, and the number and dispersion of the glowing ommatidia was increased. However, when dopaminergic Rab10 is depleted in the *THG2* deGradFP fly, the DPP is much less disrupted (*YFP-Rab10*; *THG2* > *vhhGFP* Fig. 3A ii-iv). There are fewer light-transmitting ommatidia and they are all adjacent. Thus, dopaminergic Rab10 reduction is sufficient to rescue the *G2019S-*induced neurodegeneration. We conclude that expression of *G2019S* leads to degeneration of the retina, including a lysosomal deficit in the pigment granules.

**Fig. 3.**
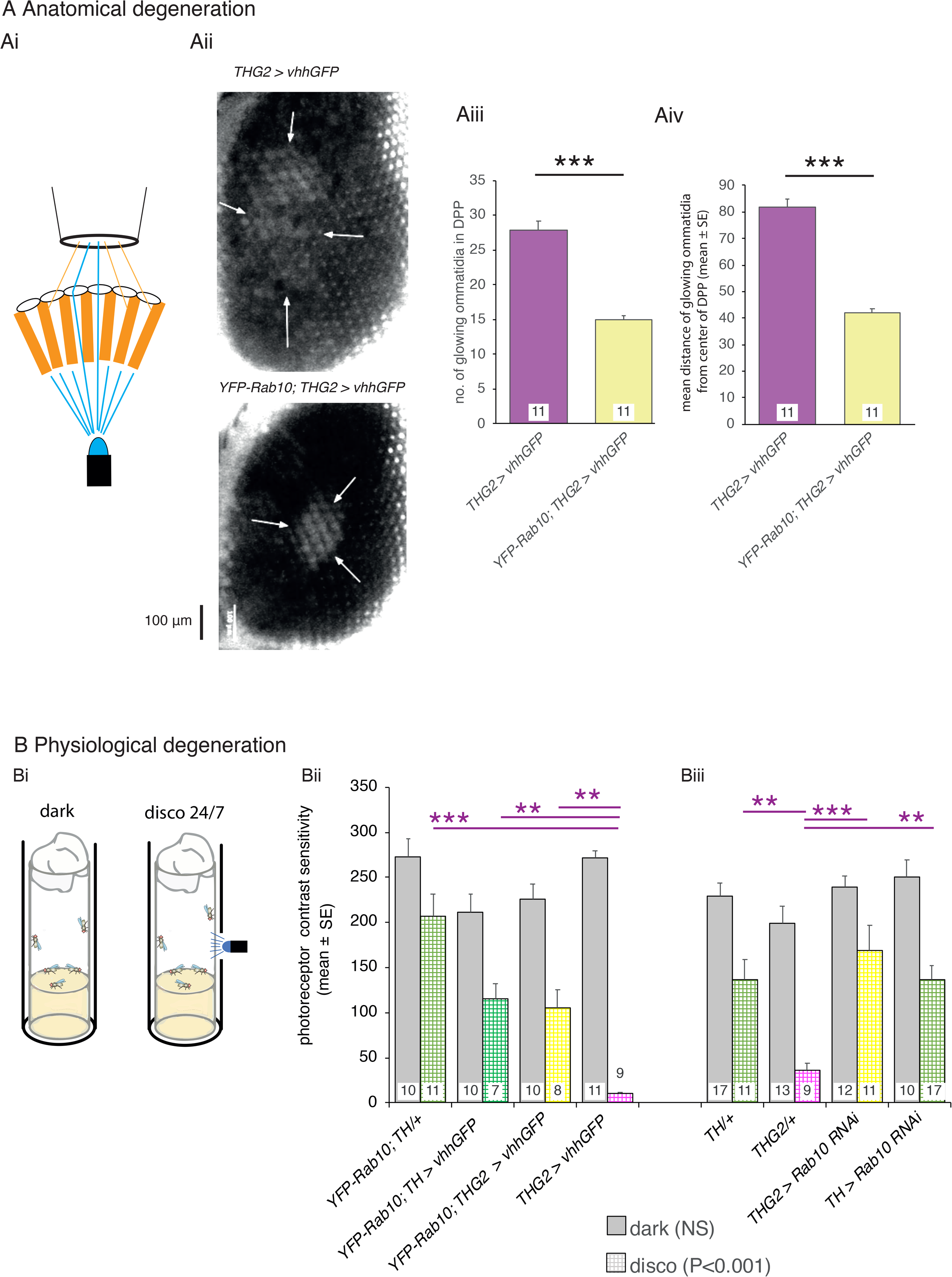
Visual degeneration due to *LRRK2-G2019S* is rescued by dopaminergic knock-down of *Rab10*. A. **Anatomically, dopaminergic *Rab10* knock-down rescues the loss of eye structure**. Ai. Healthy flies have a marked DPP (Deep-Pseudo-Pupil) which can be seen when the eye is illuminated from underneath. Light passing directly though the eye is focused towards the observer, but light at an angle is blocked by the pigmentation in the ommatidia. Aii-Aiv. PD-mimic flies (*THG2*; *vhhGFP*) kept in the dark for 14 days lose their focused DPP as the eyes show an increased number of ommatidia transmitting light, spread over a wider area, with the loss of pigmentation indicating lysosomal dysfunction. Much less degeneration is seen when Rab10 is depleted using the deGradFP technique (*YFP*-*Rab10; THG2*; *vhhGFP* flies). B. **Physiologically, dopaminergic *Rab10* knock-down rescues neuronal vision.** Bi. Degeneration is accelerated by a mild visual stress achieved by keeping the flies in ‘disco-chambers’ where the light is turned on and off approximately every 2 seconds; these flies were compared with those kept in the dark. Bii-iii. Flies kept in the dark have normal visual responses after 7 days. However, *THG2* flies in disco chambers lose their physiological response after 7 days (hatched magenta bar, controls in green). This is rescued by deGradFP mediated knockdown of Rab10 protein (Bi, *Rab10 GFP; THG2; vhhGFP* hatched yellow bar) or by dopaminergic knock-down of Rab10 by *RNAi* (Bii, *THG2* > *Rab10RNAi* yellow bar). Exact genotypes and statistical data in Table S3.

In People with Parkinson’s, the retinal degeneration of dopaminergic amacrine cells is accompanied by changes in contrast detection e.g. [33]. We therefore tested if the anatomical degeneration of the fly eye is linked to a loss of visual physiological contrast sensitivity. To do this, we used the SSVEP (Steady State Visual Evoked Potential) method which is automated to distinguish the photoreceptor contrast sensitivity from the synaptic response of the second order lamina neurons. At 7 days, all dark-reared control and *THG2* flies show a strong SSVEP photoreceptor signal, indicating that the eyes are functioning normally. There is no effect of the manipulation of *Rab10* (Fig. 3B, solid bars). On the other hand, when exposed to a mild visual stress to accelerate neurodegeneration (by being kept in the disco chamber for 1 week [15]), the *THG2* flies lose almost all their visual response – it is reduced to 18 % of the wild-type dark level (Fig. 2Bii,iii, hatched magenta bars; controls in green), indicating loss of photoreceptor function. As with the anatomical measure, the contrast sensitivity of the eye was rescued using the deGradFP manipulation (*YFP-Rab10*; *THG2* > *vhhGFP*, Fig. 2Bi, yellow bar). With co-expression of *Rab10*^*RNAi*^, this *THG2* deficit is also fully rescued (Fig. 3Dii, yellow bar).

Thus, both electrophysiological response and structural observation of the DPP indicate that *Rab10* knock-down rescues the effect of dopaminergic *LRRK2-G2019S*.

### LRRK2-induced daily activity deficits are not affected by Rab10

A third behavior modulated by dopaminergic neurons is the daily sleep-wake rhythm, both in human [34] and fly [35] (see [36] for review). The dopaminergic neurons in the fly mushroom bodies are key players in the daily activity rhythm. Locomotor activity during light /dark cycles provides a measure of circadian rhythm and sleep-wake behavior (Fig. 4A). Periods of inactivity longer than 5 minutes defined as ‘sleep’ (see [37]). In 12h:12h light on:off cycles, all flies tested show a daily sleep-wake rhythm, sleeping more in the light than in the dark, with *THG2* flies sleeping ~15% more than the controls during the ‘day’ (Fig. 4B). At the light / dark transition, all flies have high, persistent activity, and little sleep. *THG2* flies continue their activity during the dark, spending less than 50% of the time ‘asleep’ in the dark that control flies managed (*TH/+*, Fig. 4B). Neither reduction nor increased expression of Rab10 in dopaminergic neurons affected the sleep / wake cycle of either the control or *LRRK2*-*G2019S* flies. Following light-dark entrainment (LD), in the dark (DD), the fly locomotor activity pattern persists with a ~24 hour cycle providing a measure of intrinsic circadian rhythms. The period of the circadian rhythm was not affected by *LRRK2*-*G2019S* or *Rab10* manipulation, though a small reduction was seen when both transgenes were expressed (Fig. 4A). This is not unexpected as the circadian period is not affected by dopamine manipulations [38].

**Fig. 4.**
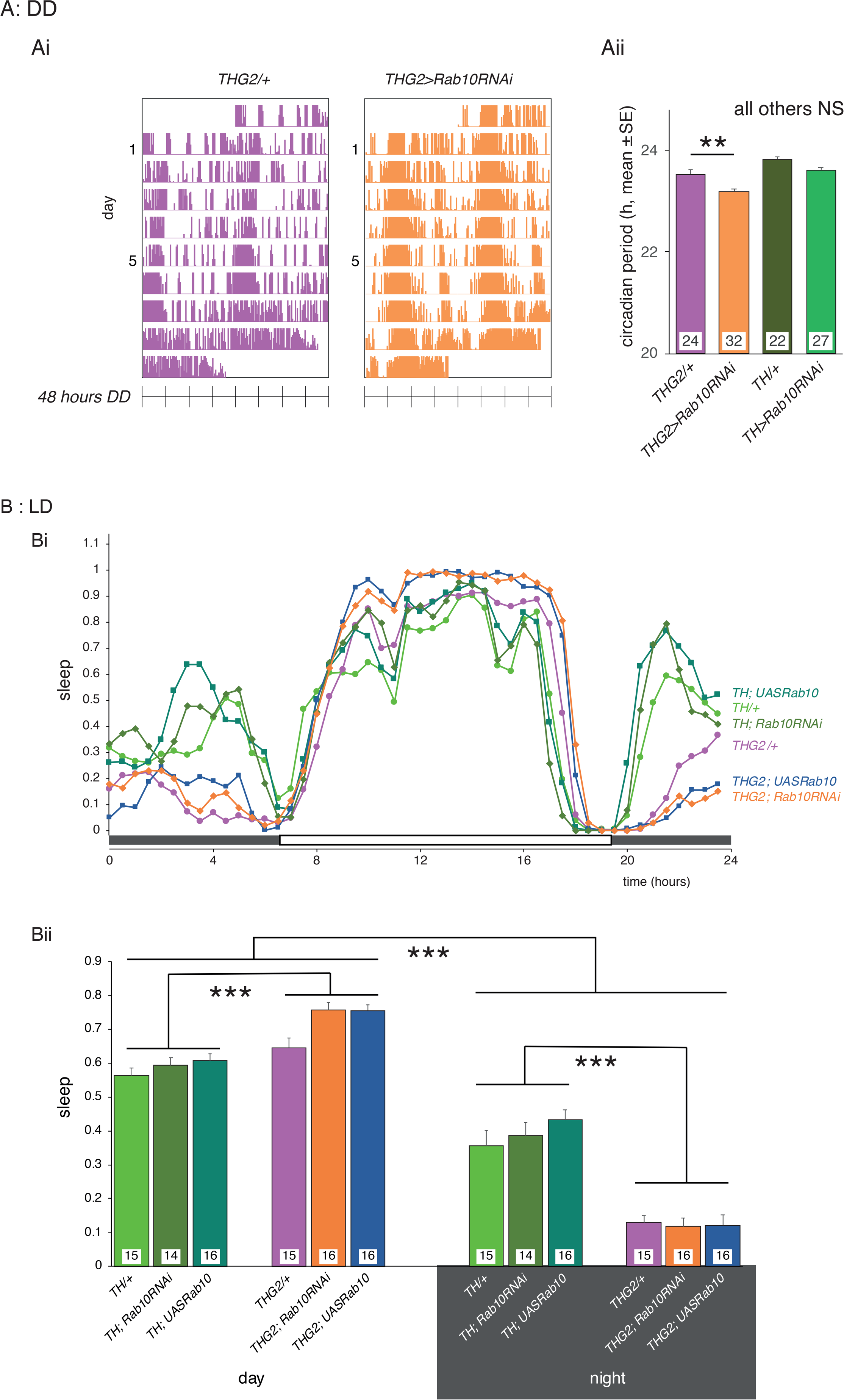
*Rab10* knock-down does not rescue sleep deficits induced by dopaminergic expression of *LRRK2-G2019S*. A. There is no effect of *LRRK2*-*G2019S* or *Rab10* expression on circadian period in continuous darkness (DD), and only a small reduction in circadian period when both genes are expressed. Ai. Raw actograms; Aii mean period from days 6-9 in DD. B. In Light /dark (LD), *THG2* flies show increased sleep during the day, and reduced sleep at night. Neither reduced nor increased expression of *Rab10* affects the daily pattern of sleep (Bi), with summary data in Bii. Sleep is defined as periods of inactivity longer than 5 min. Exact genotypes and statistical results in Table S5.

Thus, the normal sleep-wake cycle is disrupted by dopaminergic *LRRK2*-*G2019S*, but not by *Rab10* manipulation, even though Rab10 in the context of LRRK2-G2019S is seen to regulate the Proboscis Extension Response and retina.

### Improved Memory due to Rab10 depletion are insensitive to LRRK2-G2019S

Dopaminergic circuits also affect memory in both human [39] and fly [40,41]. We deployed the conditioned courtship short-term memory protocol, which is dependent on the well-known dopaminergic MB neurons [42] to assess the effects of Rab10 and *G2019S*. In this assay, males are allowed to court mated females, which reject their advances. When presented with more receptive virgin females, they tend to court less as they ‘remember’ their rejection. We found that dopaminergic knock-down of Rab10 substantially reduced the memory index, with over half of the *TH > Rab10*^*RNAi*^ flies having a memory index less than 0.05, i.e. they have excellent memory. In contrast, only 10% of control flies had a memory index less than 0.05 (Fig. 5A). When *G2019S* was expressed, no change in the distributions was seen, and Rab10 depletion still improved memory (Fig. 5B).

**Fig. 5.**
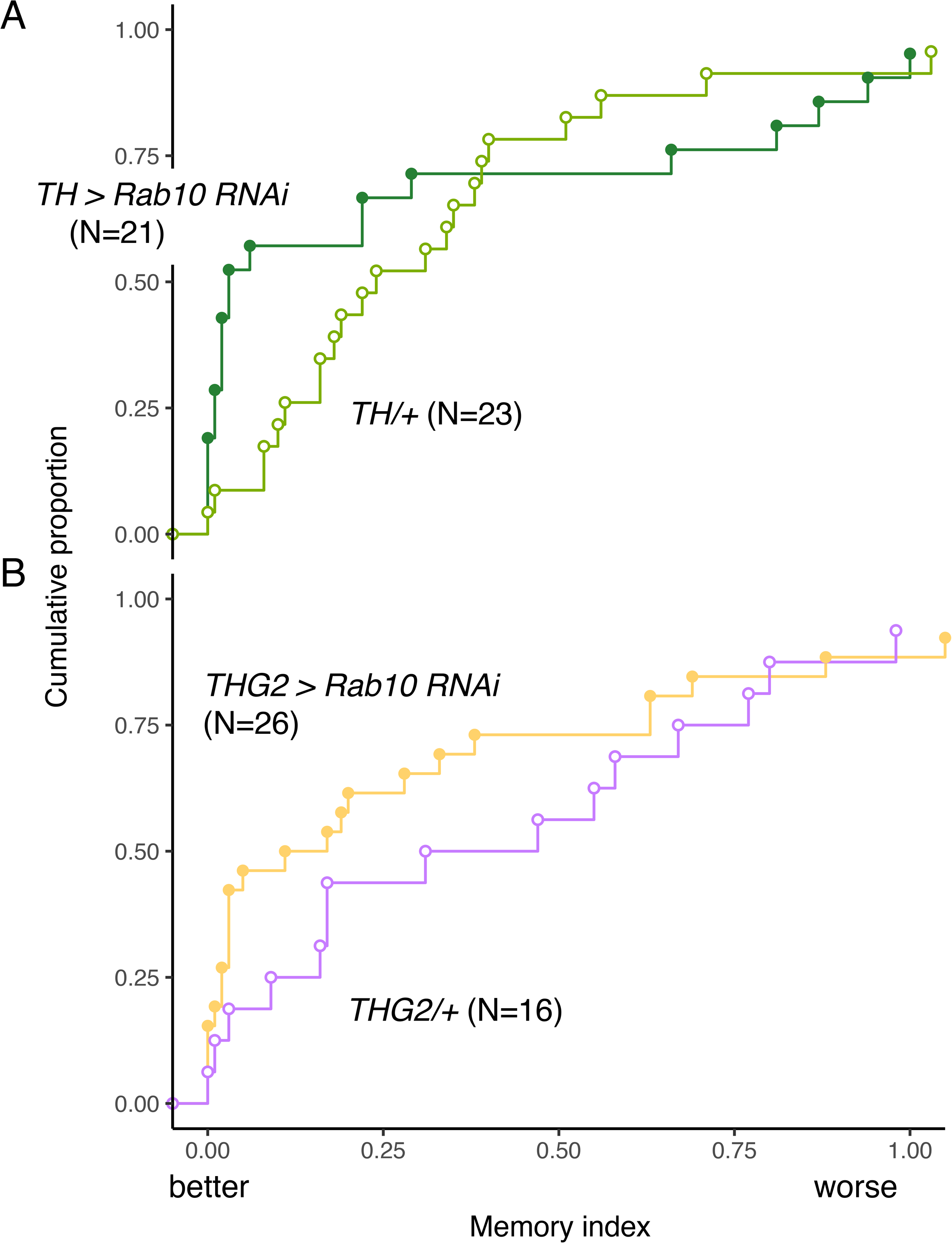
Memory depends on dopaminergic *Rab10* but not *LRRK2*-*G2019S*. A. Depletion of Rab10 in the control background (*TH* > *Rab10*^*RNAi*^) increases the proportion of flies with low memory index (MI) compared to flies with no transgene expression (*TH* / +). B. Expression of *LRRK2*-*G2019S* has no effect on performance; either for the *THG2* / + v *TH* /+ control flies, nor for the flies with *Rab10*^*RNAi*^. Exact genotypes and statistical results in Table S6.

We conclude that, in the conditioned courtship assay, Rab10 is present and has a key role in the dopaminergic neurons, but that the behavior is not affected by dopaminergic expression of *G2019S*.

### Cytoplasmic Location of G2019S and Rab10

In *THG2* flies, ectopically expressed LRRK2-G2019S protein is detected in dopaminergic neurons more strongly in the cytoplasm than the nucleus (Fig. 6A). The highest expression intensity is seen in the cytoplasm surrounding the nucleus and is quite uneven, with small holes, possibly representing the absence of the LRRK2 protein from lysosomes or mitochondria. Additionally, there is very weak staining along the axons and of the synaptic endings, including the mushroom bodies and fan-shaped body (Fig. 6C). A similar pattern of staining is seen with dopaminergic expression of *hLRRK2-wild-type* (data not shown). Rab 10 expression (*TH > Rab10-YFP* flies, Fig. 6B) shows a staining in the cytoplasm, with the strongest fluorescence at the synaptic terminals, noticeably in the neuropil of the mushroom bodies and fan-shaped body.

**Fig. 6.**
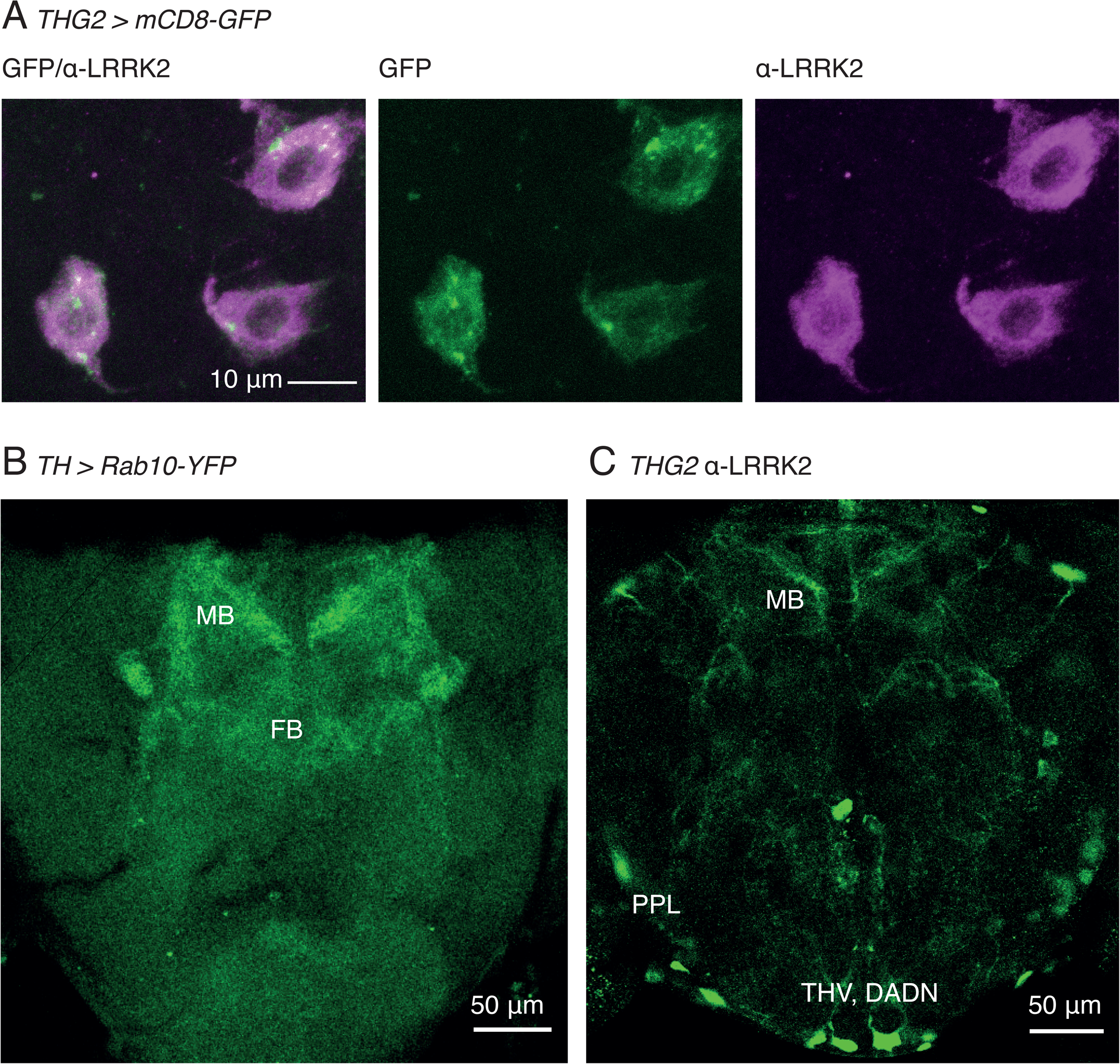
LRRK2-G2019S is found in the cytoplasm of the cell soma and in the synaptic endings of dopaminergic neurons. A. Confocal stack through the TH-VUM and DADN neurons in the ventral part of the brain of a *THG2* > *mCD8*-GFP fly. Note that LRRK2 protein is found more in the cytoplasm than the nucleus, and that there are areas of the cytoplasm with less LRRK2 staining. *mCD8*-GFP expression is used to mark dopaminergic neurons. B. Dopaminergically expressed Rab10-YFP is found in the synaptic endings in the mushroom bodies (MB) and weakly in the fan-shaped body (FB). Stack of two confocal images. C. *THG2* flies show LRRK2 protein in the dopaminergic cell bodies (PPL, THV, DADN) and in the synaptic neuropil, of the mushroom bodies. Single confocal section with α-LRRK2 antibody. Images representative of at least 3 preparations. Exact genotypes in Table S7.

## Discussion

Having previously demonstrated a strong genetic interaction, *in vivo,* between LRRK2 and Rab10 [9], we sought to determine whether manipulation of Rab10 affected the physiological deficits associated with flies expressing LRRK2 G2019S. We have identified a requirement for Rab10 in mediating the movement and visual deficits induced by dopaminergic expression of *LRRK2*-*G2019S*. In contrast, in two other dopaminergic physiological systems, there was no interaction: in one (the circadian sleep / wake cycle) *LRRK2*-*G2019S* has a marked phenotype independent of Rab10; in the other (conditioned courtship memory) Rab10 has a distinct role, but there is no *LRRK2*-*G2019S* phenotype. As such, we have begun to identify the individual dopaminergic circuits in the fly that are most sensitive to LRRK2-Rab10 interplay.

### Rab10 knock-down rescues movement and visual deficits from dopaminergic LRRK2-G2019S expression

Movement and visual deficits induced by LRRK2-G2019S expression have been linked to low dopamine release in the TH-VUM and some PPL /MC neurons that control the PER and vision respectively [14,15]. We focus here on how the LRRK2-Rab10 interaction might lead to a potential loss of dopamine release. First, *Rab10* is strongly expressed in these neurons [9] and so could play a role in controlling dopamine traffic or exocytosis. *LRRK2*-*G2019S* expression in the fly CNS increases the phosphorylation of Rab10 approximately two-fold. Knock-down by *Rab10*^*RNAi*^ reduces the level of pRab10 below the level at which it is detected, though some Rab10 is still present. Additionally, protein knock-down using deGradFP lowers pRab10 and rescues the proboscis extension /visual deficits, though the reduction in Rab10 reported by Western Blot was not so strong as with *Rab10*^*RNAi*^. Thus, the amelioration, *in vivo* of *G2019S*-induced movement and visual deficits by *Rab10*^*RNAi*^ is in accord with data from cell culture that Rab10 is a key target of LRRK2 [2–7]. Our data additionally shows, at least for a subset of Drosophila dopaminergic neurons, Rab10 is likely the relevant physiological substrate for LRRK2-G2019S.

This genetic interaction between Rab10 and LRRK2-G2019S does not imply that the physical interaction is direct, though the anatomical evidence shows that both proteins spread along the axons to the synaptic endings and are likely in proximity. Rab proteins often work in functionally linked chains e.g. [43,44]. Our data does not exclude that the *LRRK2*-*G2019S* deficits also involve other Rabs.

The cellular consequences of phosphorylation of Rab10 are not yet fully clear, but it has been suggested that this may switch the effectors to which Rab10 binds [8,44,45]. In dividing cells in culture phosphorylation of Rab10 by LRRK2 reduces ciliogenesis [46,47]. However, the relevance of this to the terminally differentiated adult Drosophila CNS neurons is not clear. Both LRRK2 and Rab10 have been linked to mitochondrial damage resolution, through the Rab10 effector OPTN (optineurin) [48] though Drosophila do not have an evident OPTN homolog (this function could be fulfilled by an as yet unidentified protein). Another role for LRRK2 – control of vesicle transport at the TGN – has support from HEK cells [49] and fly neurons. In the fly retina, a Rab10 interaction has been reported occurring at the TGN with its GEF Crag and effector Ehbp1 [50], affecting vesicle budding. Interestingly, the fly ortholog of LRRK2, which is dLRRK, is linked to the golgin protein, Lava lamp, at Golgi outposts in neurites [51]. An effect on vesicle transport at the TGN, along with consequent endosomal disruption [43,52–54] may provide an explanation for a lack of dopamine release.

### Sleep/wake cycles and conditioned courtship memory do not depend on an interaction between Rab10 and LRRK2-G2019S

In stark contrast, in the same animals, dysfunction in other dopamine-mediated behaviours does not require Rab10. *Rab10*^*RNAi*^ did not ameliorate the change in daily sleep defect induced by LRRK2-G2019S, and overexpression of Rab10 did not phenocopy G2019S-induced sleep defects. While *Rab10*^*RNAi*^ expression affected the conditioned courtship memory, indicating the presence of Rab10 and its importance in these dopaminergic neurons, *LRRK2*-*G2019S* had no phenotype. Thus, Rab10 seems to be present, but not affected by *LRRK2*-*G2019S*.

Dopaminergic neurons display varied levels of Rab10 even within a particular cluster [9]. Consequently, it is possible that the Rab10 GEFs, GAPs and effectors may also differ between the subsets of the dopaminergic neurons in the PAM, PPL1 and PPM3 neurons, the clusters that regulate the sleep-wake cycle [35,55,56]. The pattern of gene expression in the three types of PAM neurons is complex [10]. The GEF *Crag* was expressed in only one type; the GAPs *Evi5* and *plx* in different types, while the effector pattern is more complex: *Ehbp1* and *Rilpl (CG11448)* in all three PAM-types but another effecto*r (Mical)* is present in none. This suggests that Rab10 function may be regulated differentially in different dopaminergic neuronal clusters. Further, *dLRRK* is expressed in the dopaminergic neurons in to optic lobe, but not in the PAM neurons [10]. The variation in gene expression may explain why some labs have identified other potential LRRK2 interactors – for example EndophilinA, in Drosophila larval glutamatergic neuromuscular synapses [57].

### Conclusion

Our key finding is that that LRRK2-G2019S may signal by several pathways, even within the dopaminergic neuron population. We conclude that an uneven distribution of Rabs, their effectors and binding partners may contribute to the differences in the rate of dopaminergic neuron degeneration in the human striatum. Multiple LRRK2 pathways may also explain why the range of Parkinson’s symptoms initiate at different ages.

## Supporting information

Sample Movie

## Acknowledgements

We are grateful for the gifts of flies from Kristin Scott, Wanli Smith, Cheng-Ting Chien, Julie Simpson, Serge Birman, and Stefan Heidmann. We also thank the York Biology Technology Facility, Bloomington Drosophila Supply Center and Flybase for their provision. Sean Sweeney, Chris MacDonald and Amy Cording kindly read a draft manuscript. We are particularly grateful to Parkinson’s UK and to their volunteers for support (K-1704, G-1804).

## Table of Abbreviations

CI: Courtship Index
DAM: Drosophila activity monitor
dLRRK: The Drosophila homolog of LRRK2
DPP: Deep-Pseudo-Pupil
FFT: Fast-Fourier Transform
Gal4 / UAS: A binary system for targeted gene expression in the fly
GAP: GTPase-activating protein
GEF: Guanine nucleotide exchange factor
LED: Light-Emitting Diode
LexA /LexOp: A second, independent binary system for targeted gene expression in the fly
LRRK2: Leucine-rich-repeat kinase2
MC: dopaminergic neurons in the optic lobe (also known as Mi15 neurons)
MI: Memory index
nSyb: neuronal Synaptobrevin
OPTN: optineurin
PAM: A cluster of dopaminergic neurons in the fly brain
PER: Proboscis Extension Response
PPL: A second cluster of dopaminergic neurons in the fly brain
PPM: Another cluster of dopaminergic neurons in the fly brain
SSVEP: Steady State Visual Evoked Potential
TGN: Trans-Golgi Network
TH: Tyrosine-Hydroxylase
TH-VUM: Tyrosine Hydoxylase Ventral Unpaired Median neuron
*THG2*: Flies with TH Gal4 recombined with UAS-LRRK2-G2019S (PD-mimic)
YFP: Yellow Fluorescent Protein

## Declarations

### Ethics approval and consent to participate

All experiments are with *Drosophila melanogaster* and do not require consent.

### Consent for publication

All authors read and approved the final manuscript.

### Availability of data and material

Full statistical and genetic data is available in the Supplementary Tables, and the raw data is available on request

### Authors’ contributions

AF, CAM, JM, CU, DJ performed experiments, SC and LW developed apparatus /methods, SC and CJHE analysed data, CJHE drafted the manuscript, which was edited by SC, JM, DJ and CJHE.

Movie M1. **Extension of the proboscis in response to an optogenetic stimulus**. A flash of blue light is used to excite the *GR5a* sugar-sensitive neurons which leads to the extension of the proboscis. The outline of the fly (red line) was determined by thresholding. The distance between the fixed reference point and the lowest point of the proboscis (cyan dots) was measured automatically on each frame. The recorded trace is shown as the green line in Fig. 2Aiii. Each frame is 6 ms. Exact genotype: *Gr5a::ReachR / +; +/+.*

